# Synthesis of three major auxins from glucose in Engineered *Escherichia coli*

**DOI:** 10.1101/347930

**Authors:** Daoyi Guo, Lihua Zhang, Sijia Kong, Zhijie Liu, Xu Chu, Xun Li, Hong Pan

**Affiliations:** Key Laboratory of Organo-Pharmaceutical Chemistry, Jiangxi Province, Gannan Normal University, Ganzhou 341000, China; Key Laboratory of Fermentation Engineering (Ministry of Education), Hubei Key Laboratory of Industrial Microbiology, Hubei Collaborative Innovation Center for Industrial Fermentation, Hubei University of Technology, Wuhan 430068, China

**Keywords:** metabolic engineering, *Escherichia coli*, auxin

## Abstract

Indole-3-acetic acid (IAA) is considered the most common and important naturally occurring auxin in plants and a major regulator of plant growth and development. In addition, phenylacetic acid (PAA) and 4-hydroxyphenylacetic acid (4HPA) can also play a role as auxin in some plants. In recent years, several microbes have been metabolically engineered to produce IAA from L-tryptophan. In this study, we showed that aminotransferase *aro8* and decarboxylase *kdc* from *Saccharomyces cerevisiae*, and aldehyde dehydrogenase *aldH* from *Escherichia coli* have broad substrate ranges and can catalyze the conversion of three kinds of aromatic amino acids (L-tryptophan, L-tyrosine or L-phenylalanine) to the corresponding IAA, 4HPA and PAA. Subsequently, three de novo biosynthetic pathways for the production of IAA, PAA and 4HPA from glucose were constructed in *E. coli* through strengthening the shikimate pathway. This study described here shows the way for the development of agricultural microorganism for biosynthesis of plant auxin and promoting plant growth in the future.

## INTRODUCTION

Indole-3-acetic acid (IAA) is the most common plant auxin and it plays a central role in modulating plant growth and development ^1^. Phenylacetic acid (PAA) and 4-Hydroxyphenylacetic acid (4HPA) can also play a role as plant auxin in some plants^2-5^. Many rhizosphere microbes can synthesize IAA which acts as a signal molecule to promote plant growth ^6,7^. IAA synthesis in microbes can regulate the growth of plants by changing the plant auxin pool^8,9^. Moreover, microbiological IAA could signifcantly increases root surface area and length, and therefore strengthen the absorption of water and nutrient by plants, which in turn stimulate the growth of microbes ^10,11^.

Multiple IAA biosynthetic pathways have been described in microbes, most of which use L-tryptophan as the starting substrate. These L-tryptophan-dependent pathways include indole-3-acetamide, indole-3-pyruvate, tryptamine, tryptophan side-chain oxidase and indole-3-acetonitrile pathways^12^. The most studied is the indole-3-pyruvic acid pathway, which involves three steps. L-tryptophan is first converted to indole-3-pyruvic acid by an aminotransferase. Subsequently, a decarboxylase enzyme converts indole-3-pyruvic acid into indole-3-acetylaldehyde. Finally, indole-3-acetylaldehyde is converted to IAA by aldehyde dehydrogenase. In the past decades, great efforts have been made in identifying the key genes of the indole-3-pyruvic acid pathway. Aminotransferases for the conversion of the L-tryptophan to indole-3-pyruvic acid has been reported in several strains^13,14^. The gene encoding indole-3-pyruvic acid decarboxylase for the conversion of the indole-3-pyruvic acid to indole-3-acetylaldehyde has been identifed in *Azospirillum brasilense*, *Enterobacter cloacae*, and *Pseudomonas putida*^15-18^. Putative indole-3-acetylaldehyde dehydrogenase genes have been identified in the genomes of *Bacillus amyloliquefaciens* FXB42, *Azospirillum brasilense* Yu62 and *Neurospora crassa* ^19-21^.

In recent years, many wild microbes have been evaluated the capacity of synthetic IAA from L-tryptophan^19,22-28^. Further, several microbes have been metabolically engineered to produce IAA from L-tryptophan ^29,30^. However, the de novo synthesis of IAA from glucose by engineered micro-organisms is lacking. In this study, we showed that aminotransferase *aro8* and decarboxylase *kdc* from *Saccharomyces cerevisiae*, and aldehyde dehydrogenase *aldH* from *Escherichia coli* have broad substrate ranges and can catalyze the conversion of three kinds of aromatic amino acids (tryptophan,tyrosine or phenylalanine) to the corresponding IAA, 4HPA and PAA. Subsequently, three de novo biosynthetic pathways for the production of IAA, PAA and 4HPA from glucose were constructed in *E. coli* through strengthening the shikimate pathway (Figure 1). This study described here shows the way for the development of agricultural microorganism for biosynthesis of plant auxin and promoting plant growth in the future.

**Figure 1.**
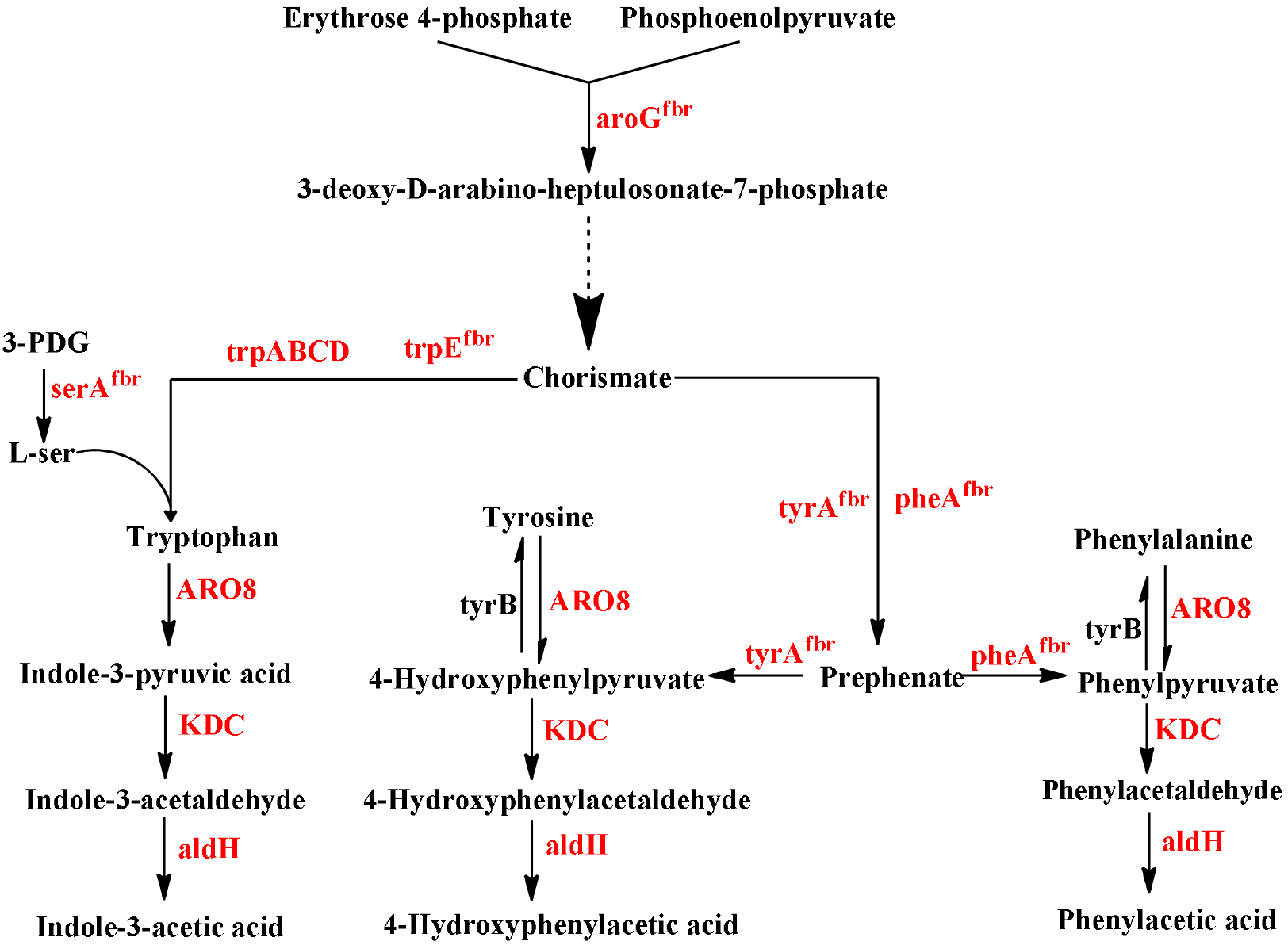
Engineered pathways for production of IAA, PAA and 4HPA from glucose.

## MATERIALS AND METHODS

### Enzymes, chemicals, and strains

Restriction endonucleases and DNA polymerases were purchased from New England Biolabs. T4 DNA ligase, plasmid extraction kits, PCR purification kits and DNA extraction kits were purchased from Fermentas (Burlington, Canada). Plasmid construction and screening were performed in *E. coli* DH1013. A reduced aromatic aldehyde reduction *E. coli* strain RARE whose seven aldehyde reductase genes (*dkgB*, *yeaE*, *yqhC*, *yqhD*, *dkgA*, *yahK*, *yjgB*) have been deleted, got the aromatic aldehydes as end products could be accumulated, was used as a host for fermentation to synthesize PAA and 4HPA. *E. coli* strain DG101 which was constructed by deleting the trpR gene using the CRISPR/Cas9 technology in strain RARE by Genewiz Biotech Co. Ltd, was used as a host for fermentation to synthesize IAA.

### Plasmid construction

Plasmid pDG11 for expression *aroG*^*fbr*^ and pDG14 for co-expression *aroG*^*fbr*^ and *pheA*^*fbr*^ gene from *E. coli* was constructed in our previous study^31^. *TyrA*^*fbr*^ (M53I/A354V, GenBank NP_417091.1) was synthesized by Genewiz Biotech Co. Ltd and amplified by PCR using primers *tyrA*^*fbr*^-*XbaI* and *tyrA*^*fbr*^-*SpeI-SalI*, and ligated into pET28a(+) via *XbaI* and *SalI* to yield plasmid pDG20. The *XbaI-SalI* fragments of *tyrA*^*fbr*^ from pDG20 was inserted into *SpeI* and *SalI* sites of pDG11 to give pDG21. The *XbaI-SalI* fragments of *aroG*^*fbr*^ and *tyrA*^*fbr*^ from pDG21 was inserted into *XbaI*- and *XhoI* sites of pBBRMCS1 to give pDG22.

*TrpE*^*fbr*^ (S40F, GenBank NP_415780.1) was synthesized by Genewiz Biotech Co. Ltd and amplified by PCR using primers *trpE*^*fbr*^ -*XbaI* and *trpE*^*fbr*^-*SpeI-BamHI* and ligated into pET28a(+) via *XbaI* and *BamHI* to yield plasmid pDG23. *SerA*^*fbr*^ (H344A/N364A, GenBank NP_417388.1) was synthesized by Genewiz Biotech Co. Ltd and amplified by PCR using primers *serA*^*fbr*^-*XbaI* and *serA*^*fbr*^-*SpeI*-*SalI* and ligated into pET28a(+) via *XbaI* and *SalI* to yield plasmid pDG24. The *XbaI-SalI* fragments of *trpE*^*fbr*^ from pDG23 was inserted into *SpeI* and *SalI* sites of pDG11 to give pDG25. The *XbaI*-*SalI* fragments of *serA*^*fbr*^ from pDG24 was inserted into *SpeI* and *SalI* sites of pDG25 to give pDG26. The *XbaI-SalI* fragments of *aroG*^*fbr*^, *tyrA*^*fbr*^ and *serA*^*fbr*^ from pDG26 was inserted into *XbaI*- and *XhoI* sites of pBBRMCS1 to give pDG27.

Plasmid pDG9 for co-expression *aro8* and *kdc* gene from *S. cerevisiae* YPH499, and *aldH* gene from *E. coli* was constructed in our previous study^32^. *TrpABCD* was amplified by PCR using primers *trpABCD*-*XbaI* and *trpABCD*-*XhoI* and ligated into pDG9 via *NheI* and *XhoI* to yield plasmid pDG28. All the primers used in this study are listed in Table 1. The bacterial strains and plasmids used in this study are listed in Table 2.

**Table 1.**
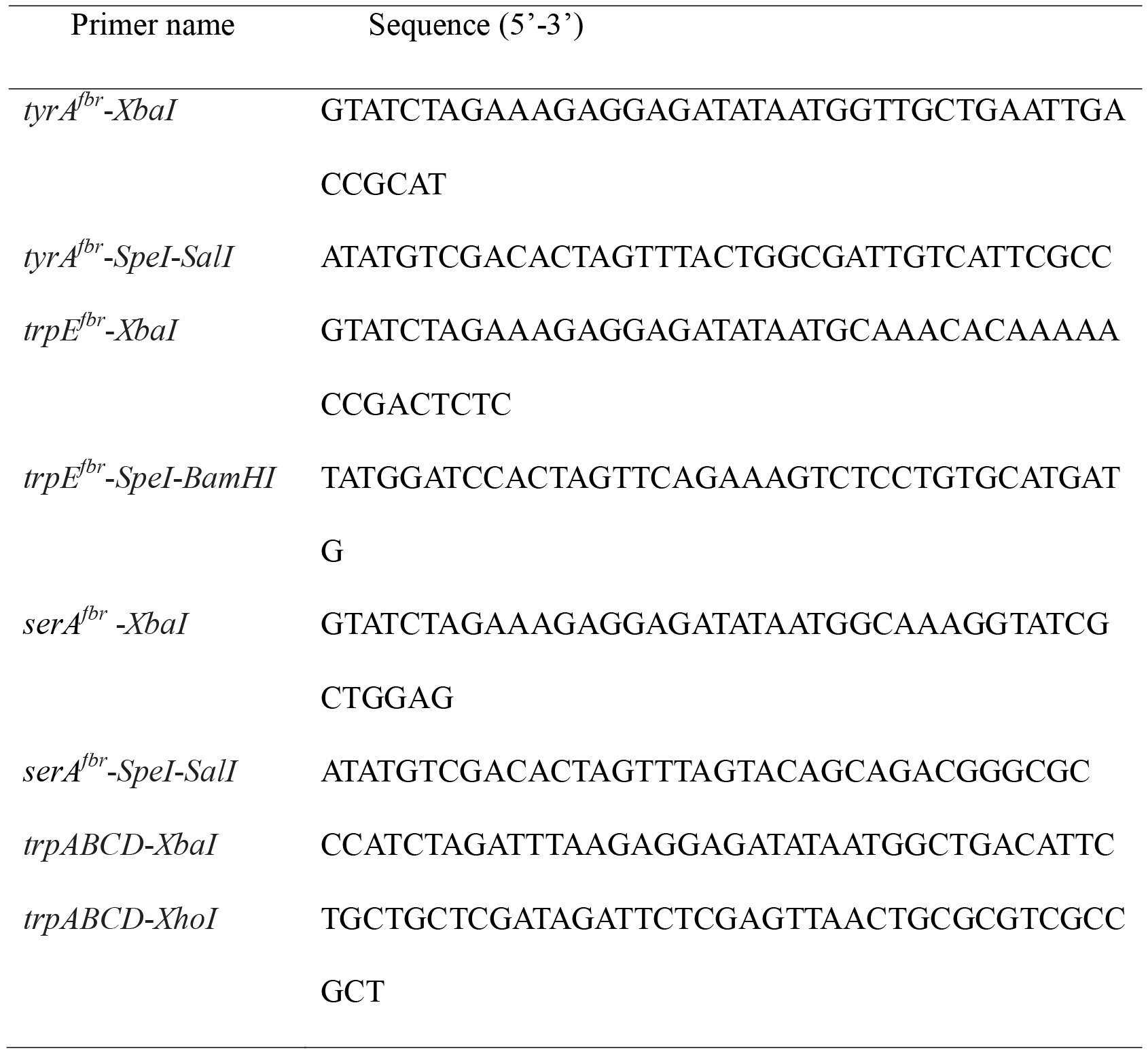
Primers used in this study

**Table 2.** Plasmids and strains used in this study

| Plasmids/strains | Properties | Source |
| --- | --- | --- |
| Plasmids |  |  |
| pDG9 | pET28a; pT7: *aro8*, *kdc* and *aldH* | 31 |
| pDG11 | pET28a; pT7: *aroG*<sup>fbr</sup> | This study |
| pDG12 | pET28a; pT7: *pheA*<sup>fbr</sup> | This study |
| pDG13 | pET28a; pT7: *aroG*<sup>fbr</sup> and *pheA*<sup>fbr</sup> | This study |
| pDG14 | pBBRMCS1; pT7: *aroG*<sup>fbr</sup> and *pheA*<sup>fbr</sup> | This study |
| pDG20 | pET28a; pT7: *tyrA*<sup>fbr</sup> | This study |
| pDG21 | pET28a; pT7: *aroG*<sup>fbr</sup> and *tyrA*<sup>fbr</sup> | This study |
| pDG22 | pBBRMCS1; pT7: *aroG*<sup>fbr</sup> and *tyrA*<sup>fbr</sup> | This study |
| pDG23 | pET28a; pT7: *trpE*<sup>fbr</sup> | This study |
| pDG24 | pET28a; pT7: *serA*<sup>fbr</sup> | This study |
| pDG25 | pET28a; pT7: *aroG*<sup>fbr</sup> and *trpE*<sup>fbr</sup> | This study |
| pDG26 | pET28a; pT7: *aroG*<sup>fbr</sup>, *trpE*<sup>fbr</sup> and *serA*<sup>fbr</sup> | This study |
| pDG27 | pBBRMCS1; pT7: *aroG*<sup>fbr</sup>, *trpE*<sup>fbr</sup> and *serA*<sup>fbr</sup> | This study |
| pDG28 | pET28a; pT7: *aro8*, *kdc*, *aldH* and *trpABCD* | This study |
| Strains |  |  |
| RARE | MG1655 derivative; $\Delta dkgB$, $\Delta yeaE$, $\Delta yqhC$, $\Delta yqhD$, $\Delta dkgA$, $\Delta yahK$ and $\Delta yjgB$ | 59 |
| DG101 | RARE derivative; $\Delta trpR$ | This study |
| DG102 | RARE/pDG9 | This study |
| DG103 | RARE/pDG9/ pDG14 | This study |
| DG104 | RARE/pDG9/ pDG22 | This study |
| DG105 | DG101/pDG27/ pDG28 | This study |

### Shake flask cultures

Experiments of shaking flask culture were performed at 30 °C in 100 mL modified M9 medium as previously described by Guo ^33^. 36 mg/L chloromycetin and 50 mg/L kanamycin were added to the medium, when needed. When the OD_600_ reached about 0.8, IPTG was added to the medium for induction of gene expression at a final concentration of 0.1 mM.

### Analytical methods

Glass beads were added to 10 mL culture broth for breaking the cells. The pH of the culture broth was adjusted to 3.0 with 3 M HCl. Ethyl acetate (15 mL) is used to extract PAA, 4HPA and IAA. After the centrifuge stratification, the upper ethyl acetate phase was withdrawn, evaporated to dryness. These extracted PAA and 4HPA was redissolved in 0.6 mL ethyl acetate. A 1 μL portion of the ethyl acetate phase was analysed after a split 10:1 injection on an Agilent 7890A GC equipped with an Agilent 5975 MS detector and an Agilent DB-5MS capillary column. The following temperature program was applied: 80 °C for 1 min, an increase of 20 °C/min to 260 °C, and 260 °C for 5 min.

When IAA is directly detected by GC/MS, IAA is partially decomposed into 3-Methylindole (Figure 2), probably due to its intolerance to high temperature gasification. To achieve better detection of IAA, IAA was converted to more stable methyl indole-3-acetate ^34^. The extracted IAA was esterified by redissolving in 1 mL 5% H_2_SO_4_ in methanol and incubating at 90 °C for 2 hr. After the reaction, 1 mL 0.9% NaCl was added into each sample, and then the methyl indole-3-acetate was extracted using 1 mL ethyl acetate for subsequent GC/MS analysis. A 1 μL portion of the n-hexane phase was analysed after a split 20:1 injection on a HP‒5MS capillary column. The following temperature program was applied: 80 °C for 1 min, an increase of 20 °C/min to 250 °C, and 250 °C for 8 min.

**Figure 2.**
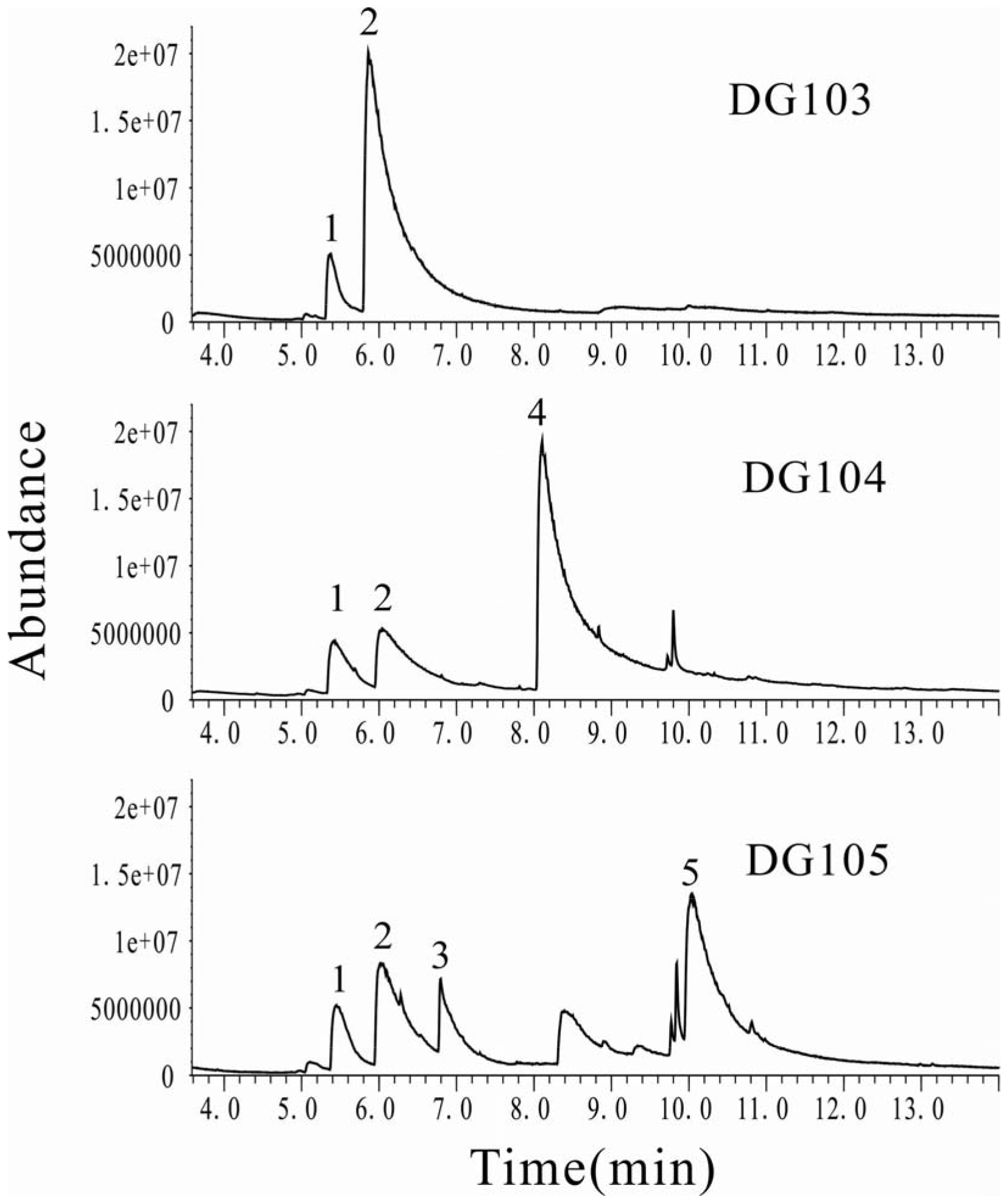
GC/MS analyses of PAA, 4HPA and IAA from glucose in engineered E. coli strains DG103, DG104 and DG105. Identified substances: 1, benzoic acid (internal standard); 2, PAA; 3, 3-Methylindole; 4, 4HPA, 5, IAA.

## RESULTS

### The biosynthesis of IAA and 4HPA from tryptophan and tyrosine in *E. coli*

Several aldehyde dehydrogenases (*feaB*, *aldB* and *aldH*) were reported to have the ability of oxidation phenylacetaldehyde or benzaldehyde in vitro ^35-37^. Recently, we have compared the activity of these aldehyde dehydrogenases for the oxidation of phenylacetaldehyde ^32^. We found that only *E.coli* AldH is an efficient aldehyde dehydrogenase for oxidation of phenylacetaldehyde to PAA in vivo. Subsequently, we showed metabolic engineering of *E. coli* to high efficient synthesis PAA from L-phenylalanine through overexpression of an aminotransferase *aro8* for the conversion of L-phenylalanine to phenylpyruvate, a keto acid decarboxylase *kdc* from *S. cerevisiae* for the decarboxylation of phenylpyruvate to phenylacetaldehyde and an aldehyde dehydrogenase *aldH* from *E.coli* for the oxidation of phenylacetaldehyde to the corresponding PAA.

To determine whether *aro8* and *kdc* from *S. cerevisiae*, and *aldH* from *E.coli* have broad substrate ranges and can catalyze the conversion of other two kinds of aromatic amino acids (L-tryptophan or L-tyrosine) to the corresponding IAA and 4HPA. *E. coli* RARE strain harboring a plasmid pDG9 for co-expression *aro8*, *kdc* and *aldH* gene was grown in modified M9 medium with 0.4 g/L of L-tryptophan (the solubility of L-tryptophan in water is about 479 mg/L) or 1 g/L L-tyrosine. The resulting *E. coli* strain DG102 produced up to 303 ± 14.1 mg/L IAA from 0.4 g/L of L-tryptophan with the molar yield of 0.91 moL/moL or 726 ± 28.3 mg/L 4HPA from 1 g/L of L-tyrosine with the molar yield of 0.86 moL/moL (Table 3). This proves that *aro8*, *kdc* and *aldH* can catalyze effectively conversion of L-tryptophan or L-tyrosine to the corresponding IAA and 4HPA.

**Table 3.**
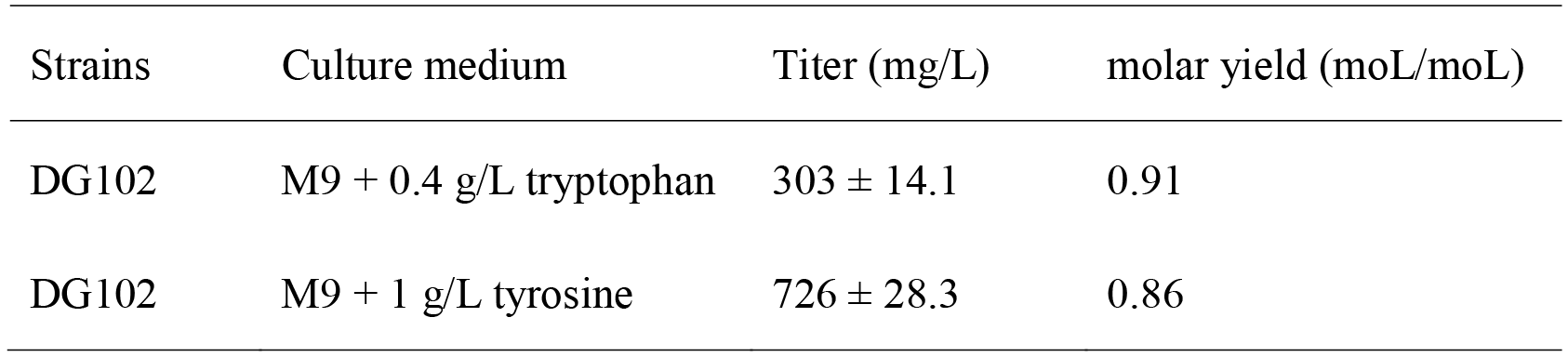
The biosynthesis of IAA and 4HPA from L-tryptophan and L-tyrosine in *E. coli* DG102. Data are the mean values of three experiments ± SD.

### Creation of a synthetic pathway for PAA in *E. coli* from glucose

The L-phenylalanine can be de novo biosynthesized from glucose via L-phenylalanine pathway. However the efficiency is quite low because of feedback inhibition in L-phenylalanine pathway. Overexpression of feedback resistant mutant of 3-deoxy-D-arabinoheptulosonate-7-phosphate synthase (*aroG*^*fbr*^) and the bifunctional enzyme chorismate mutase/prephenate dehydratase (*pheA*^*fbr*^) has proven to be an effective strategy to overproduce L-phenylalanine from glucose ^38-40^. In this study, a PAA artificial biosynthesis pathway was constructed in *E. coli* strain RARE by co-expression of two feedback-resistant mutants *aroG*^*fbr*^ and *pheA*^*fbr*^ for the efficient overproduction of L-phenylalanine, aminotransferase *aro8*, decarboxylase *kdc* and aldehyde dehydrogenase *aldH* for conversation of L-phenylalanine to PAA. The resulting *E. coli* strain DG103 was cultured in modified M9 medium with 20 g/L of glucose and produced up to 1217 ± 45.2 mg/L PAA within 28 h (Figure 2). PAA yield and productivity were calculated to be 60.8 mg/g glucose and 43.7 mg/L/h, respectively (Table 4). This proves that the designed PAA synthetic pathway was functionally expressed in *E. coli*.

**Table 4.** Titer, yield and productivity of PAA, 4HPA and IAA by engineered *E. coli* strains in shake flasks for 28 h. Yield calculated by milligrams PAA, 4HPA or IAA produced divided by grams glucose consumed. Productivity calculated by concentration of PAA, 4HPA or IAA produced per hour. Data are the mean values of three experiments ± SD.

| Strains | product | Titer (mg/L) | Yield (mg/g) | Productivity (mg/L/h) |
| --- | --- | --- | --- | --- |
| DG103 | PAA | 1217 ± 45.2 | 60.8 ± 2.26 | 43.7 ± 1.62 |
| DG104 | 4HPA | 991 ± 48.8 | 49.5 ± 2.44 | 35.4 ± 1.74 |
| DG105 | IAA | 1265 ± 87.8 | 63.3 ± 4.38 | 45.2 ± 3.13 |

### Creation of a synthetic pathway for 4HPA from glucose

After successfully demonstrating the biosynthesis of 4HPA from L-tyrosine in *E. coli*, we further designed the biosynthesis of 4HPA from glucose. The L-tyrosine can be de novo biosynthesized from glucose via L-tyrosine pathway ^41,42^. Overexpression of feedback resistant mutant of *aroG*^*fbr*^ and chorismate mutase/prephenate dehydrogenase *tyrA*^*fbr*^ is an essential strategy to overproduce L-tyrosine ^43,44^. In this study, a 4HPA biosynthesis pathway was constructed in *E. coli* strain RARE by co-expression of two feedback-resistant mutants (*aroG*^*fbr*^ and *tyrA*^*fbr*^) for the efficient overproduction of L-tyrosine, *aro8*, *kdc* and *aldH* for conversation of L-tyrosine to 4HPA. The resulting *E. coli* strain DG104 produced up to 991 ± 48.8 mg/L 4HPA within 28 h (Figure 2). 4HPA yield and productivity were calculated to be 49.5 mg/g glucose and 35.4 mg/L/h, respectively (Table 4). This proves that the designed 4HPA synthetic pathway was functionally expressed in *E. coli.*

### Creation of a synthetic pathway for IAA from glucose

After successfully confirming the biosynthesis of IAA from L-tryptophan in *E. coli*, we further designed the biosynthesis of IAA from glucose. To obtain IAA from glucose, candidate strains need have a strong capacity for the L-tryptophan synthesis. Recently, great efforts have been made on engineered *E. coli* with the goal of efficient production of L-tryptophan from glucose ^45,46^. Indole and L-serine are two precursor substrates for the biosynthesis of L-tryptophan. Therefore, increasing the availability of indole and L-serine is requisite for efficient biosynthesis of L-tryptophan.

In order to overproduce precursor substrate L-tryptophan, several strategies have been implemented in this study, such as knocking out L-tryptophan transcriptional repressor trpR to eliminate transcription regulation of the genes in L-tryptophan pathway, overexpression of feedback resistant derivatives of *aroG*^*fbr*^ and anthranilate synthase *trpE*^*fbr*^ to increase the carbon flow from glucose to indole, overexpression of feedback resistant derivatives of *serA*^*fbr*^ to increase the availability of L-serine, and overexpression of *trpABCD* for the efficient synthesis of L-tryptophan from indole and L-serine. Subsequently, a complete IAA biosynthesis pathway was established by further co-expression of *aro8*, *kdc* and *aldH* for conversation of L-tryptophan to IAA. The resulting *E. coli* strain DG105 produced up to 1265 ± 87.8 mg/L IAA within 28 h (Figure 3). IAA yield and productivity were calculated to be 63.3 mg/g glucose and 45.2 mg/L/h, respectively (Table 4). This proves that the designed IAA synthetic pathway from glucose was functionally expressed in *E. coli*.

**Figure 3.**
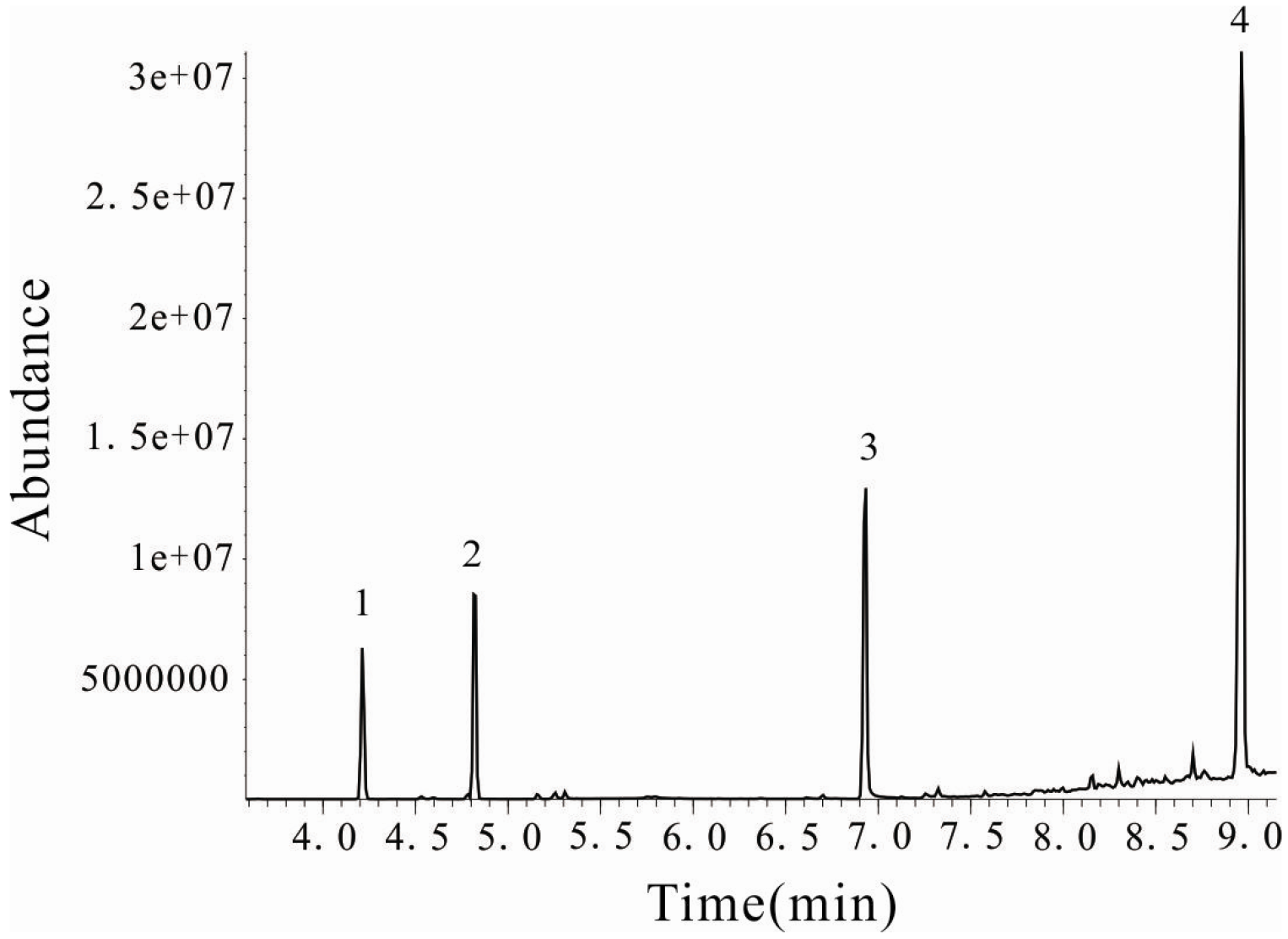
The extracted IAA from engineered *E. coli* strains DG105 was esterified and detected by GC/MS. Identified substances: 1, methyl benzoate (internal standard); 2, methyl phenylacetate: 3, methyl 4-hydroxyphenylacetate; 4, methyl indole-3-acetate.

## DISCUSSION

Recently, great efforts have been made in developing sustainable methods that could replace traditional chemical synthesis to produce valuable compounds ^47-50^. Based on synthetic biology technology, varieties of engineered microbes have been made to produce fuels, bulk chemicals and pharmaceuticals ^51-55^. The production of aromatic molecules which derived from the common shikimate pathway is a key research area and has attracted a great deal of attention in recent years ^56-59^. In this pathway, two glycometabolic intermediates erythrose 4-phosphate and phosphoenolpyruvate are first assembled into 3-deoxy-D-arabino-heptulosonate-7-phosphate (DAHP), which is then converted to chorismate. Subsequently, chorismate can be easily converted into a variety of aromatic molecules via a limited number of steps ^35^. These aromatic molecules are widely used in the paper and dye manufacture, pharmaceuticals, cosmetics, and food industries.

The large-scale use of chemical fertilizers and pesticides has caused serious environmental pollution. Great efforts have been made on the use of microorganisms instead of chemical fertilizers and pesticides to support plant growth through direct and indirect mechanisms. Among these beneficial microbes, the auxin-producing strain has played an important role in promoting plant growth. Aromatic molecules IAA, PAA and 4HPA are three major types of auxin in plants. By improving the levels of auxin produced by microbe through synthetic biology technology, it is possible to develop an agricultural microorganism to promote plant growth. Great efforts have been made on engineered microbe for the biosynthesis of auxin from L-tryptophan. Shao et al. reported co-expression of the aminotransferase *patB*, decarboxylase *yclC*, indole-3-acetaldehyde dehydrogenase *dhaS* in *Bacillus amyloliquefaciens* SQR9 and *Bacillus subtilis* 168 produced up to 39 mg/L and 22 mg/L of IAA from 3 mM tryptophan, respectively. Overexpression of putative IAA biosynthesis genes from *Pseudomonas sp* UW4 in the wild-type strain resulted in a significant increase of IAA production from tryptophan^30^. However, compared with L-tryptophan, cheaper substrate should be used to reduce the cost of production of auxin.

In this study, three de novo biosynthetic pathways for the production of IAA, PAA and 4HPA from glucose were constructed in *E. coli* through strengthening the shikimate pathway. The overall pathway includes the upstream pathway from glucose to aromatic amino acids and the downstream pathway from aromatic amino acids to auxin. Microorganic synthesis of auxin is impeded by the formation of by-product aromatic alcohols due to quick and endogenous conversion of aromatic aldehydes to their corresponding aromatic alcohols in *E. coli.* Kunjapur constructed an *E. coli* strain RARE that can accumulate aromatic aldehydes as end products by deleting seven aldehyde reductase genes ^60^. In this study, *E. coli* strain RARE was used as the host to prompt more PAA, 4HPA and IAA biosynthesis.

## CONCLUSION

In summary, we report the discovery of *S. cerevisiae* aminotransferase *aro8*, decarboxylase *kdc* and *E. coli* aldehyde dehydrogenase *aldH* from have broad substrate ranges. We successfully generated engineered *E. coli* strains capable of producing three main kinds of auxins from glucose and showed the way for the development of agricultural microorganism for biosynthesis of auxin and promoting plant growth in the future.

## Acknowledgments

This work was supported by National Natural Science Foundation of China (81460312).

## References

(1) Bartel, B. Annual Review of Plant Physiology & Plant Molecular Biology 1997, 48, 51.

(2) Pearse, H. L. Nature 1936, 138, 363.

(3) Johnson, C. F.; Morris, D. A. Planta 1987, 172, 400.

(4) Fries, L.; Iwasaki, H. Plant Science Letters 1976, 6, 299.

(5) Sugawara, S.; Mashiguchi, K.; Tanaka, K.; Hishiyama, S.; Sakai, T.; Hanada, K.; Kinoshita-Tsujimura, K.; Yu, H.; Dai, X.; Takebayashi, Y. Plant & Cell Physiology 2015, 56, 1641.

(6) Spaepen, S.; Vanderleyden, J.; Remans, R. Fems Microbiology Reviews 2007, 31, 425.

(7) Li, M.; Guo, R.; Yu, F.; Chen, X.; Zhao, H.; Li, H.; Wu, J. International Journal of Molecular Sciences 2018, 19, 443.

(8) Dobbelaere, S.; Croonenborghs, A.; Thys, A.; Broek, A. V.; Vanderleyden, J. Plant & Soil 1999, 212, 155.

(9) Ahemad, M.; Kibret, M. Journal of King Saud University-Science 2014, 26, 1.

(10) Ahmed, A.; Hasnain, S.; Bai, F. W.; Zhang, W.; Zhong, J. J. Pure & Applied Chemistry 2010, 82, 313.

(11) Parray, J. A.; Jan, S.; Kamili, A. N.; Qadri, R. A.; Egamberdieva, D.; Ahmad, P. Journal of Plant Growth Regulation 2016, 35, 877.

(12) Duca, D.; Lorv, J.; Patten, C. L.; Rose, D.; Glick, B. R. Antonie Van Leeuwenhoek 2014, 106, 85.

(13) Kittell, B. L.; Helinski, D. R.; Ditta, G. S. Journal of Bacteriology 1989, 171, 5458.

(14) Koga, J.; Syono, K.; Ichikawa, T.; Adachi, T. Biochim Biophys Acta 1994, 1209, 241.

(15) Cl, P.; Br, G. Applied and Environmental Microbiology 2002, 68, 3795.

(16) Baudoin, E.; Lerner, A.; Mirza, M. S.; Elzemrany, H.; Prigentcombaret, C.; Jurkevich, E.; Spaepen, S.; Vanderleyden, J.; Nazaret, S.; Okon, Y. Research in Microbiology 2010, 161, 219.

(17) Costacurta, A.; Keijers, V.; Vanderleyden, J. Molecular & General Genetics Mgg 1994, 243, 463.

(18) Koga, J.; Adachi, T.; Hidaka, H. Mol Gen Genet 1991, 226, 10.

(19) Idris, E. E.; Iglesias, D. J.; Talon, M.; Borriss, R. Molecular plant-microbe interactions: MPMI 2007, 20, 619.

(20) Xie, B.; Xu, K.; Zhao, H. X.; Chen, S. F. Fems Microbiology Letters 2010, 248, 57.

(21) Sardar, P.; Kempken, F. Plos One 2018, 13, e0192293.

(22) Apine, O. A.; Jadhav, J. P. Journal of Applied Microbiology 2011, 110, 1235.

(23) Kochar, M.; Upadhyay, A.; Srivastava, S. Research in Microbiology 2011, 162, 426.

(24) Tsavkelova, E. A.; Cherdyntseva, T. A.; Botina, S. G.; Netrusov, A. I. Microbiological Research 2007, 162, 69.

(25) Mauricio, S.; Claudiam, R.; Martina, P.; Marial, C.; Luigi, C.; Josea, C. Soil Biology & Biochemistry 2009, 41, 1768.

(26) Jasim, B.; Jimtha, J. C.; Shimil, V.; Jyothis, M.; Radhakrishnan, E. K. Journal of Applied Microbiology 2015, 117, 786.

(27) Nutaratat, P.; Monprasit, A.; Srisuk, N. Biotech 2017, 7, 305.

(28) Ghosh, P. K.; Sen, S. K.; Maiti, T. K. Biocatalysis & Agricultural Biotechnology 2015, 4, 296.

(29) Shao, J.; Li, S.; Zhang, N.; Cui, X.; Zhou, X.; Zhang, G.; Shen, Q.; Zhang, R. Microbial Cell Factories 2015, 14, 130.

(30) Duca, D. R.; Rose, D. R.; Glick, B. R. Antonie Van Leeuwenhoek 2018, 1.

(31) Guo, D.; Zhang, L.; Kong, S.; Liu, Z.; Li, X.; Pan, H. Journal of Agricultural and Food Chemistry 2018.

(32) Zhang, L.; Liu, Q.; Pan, H.; Li, X.; Guo, D. Amb Express 2017, 7, 105.

(33) Guo, D.; Zhu, J.; Deng, Z.; Liu, T. Metabolic Engineering 2014, 22, 69.

(34) Voelker; A. T.; Davies; M. H. Journal of Bacteriology 1994, 176, 7320.

(35) Koma, D.; Yamanaka, H.; Moriyoshi, K.; Ohmoto, T.; Sakai, K. Applied & Environmental Microbiology 2012, 78, 6203.

(36) Jo, J. E.; Mohan, S. R.; Rathnasingh, C.; Selvakumar, E.; Jung, W. C.; Park, S. Applied Microbiology & Biotechnology 2008, 81, 51.

(37) Ho, K. K.; Weiner, H. Journal of Bacteriology 2005, 187, 1067.

(38) Báez-Viveros, J. L.; Osuna, J.; Hernández-Chávez, G.; Soberán, X.; Bolívar, F.; Gosset, G. Biotechnology & Bioengineering 2004, 87, 516.

(39) Liu, S. P.; Xiao, M. R.; Zhang, L.; Xu, J.; Ding, Z. Y.; Gu, Z. H.; Shi, G. Y. Process Biochemistry 2013, 48, 413.

(40) Liu, S. P.; Liu, R. X.; Xiao, M. R.; Zhang, L.; Ding, Z. Y.; Gu, Z. H.; Shi, G. Y. Process Biochemistry 2014, 49, 751.

(41) Santos, C. N.; Xiao, W.; Stephanopoulos, G. Proceedings of the National Academy of Sciences of the United States of America 2012, 109, 13538.

(42) Santos, C. N. S.; Xiao, W.; Stephanopoulos, G. Proceedings of the National Academy of Sciences of the United States of America 2012, 109, 13538.

(43) Juminaga, D.; Baidoo, E. E. K.; Reddingjohanson, A. M.; Batth, T. S.; Burd, H.; Mukhopadhyay, A.; Petzold, C. J.; Keasling, J. D. Applied & Environmental Microbiology 2012, 78, 89.

(44) Santos, C. N. S.; Stephanopoulos, G. Applied & Environmental Microbiology 2008, 74, 1190.

(45) Chen, L.; Zeng, A. P. Appl Microbiol Biotechnol 2017, 101, 559.

(46) Gu, P.; Yang, F.; Li, F.; Liang, Q.; Qi, Q. Appl Microbiol Biotechnol 2013, 97, 6677.

(47) Rodriguez, G. M.; Tashiro, Y.; Atsumi, S. Nature Chemical Biology 2014, 10, 259.

(48) Beller, H. R.; Rodrigues, A. V.; Zargar, K.; Wu, Y. W.; Saini, A. K.; Saville, R. M.; Pereira, J. H.; Adams, P. D.; Tringe, S. G.; Petzold, C. J. Nature Chemical Biology 2018.

(49) Akhtar, M. K.; Turner, N. J.; Jones, P. R. Prtas 2013, 110, 87.

(50) Gupta, A.; Reizman, I. M. B.; Reisch, C. R.; Prather, K. L. J. Nature Biotechnology 2017, 35, 273.

(51) Steen, E. J.; Kang, Y. S.; Bokinsky, G.; Hu, Z. H.; Schirmer, A.; Mcclure, A.; Cardayre, S. B. D.; Keasling, J. D. Nature 2010, 463, 559.

(52) Atsumi, S.; Hanai, T.; Liao, J. C. Nature 2008, 451, 86.

(53) Kalscheuer, R.; Stölting, T.; Steinbüchel, A. Microbiology 2006, 152, 2529.

(54) Avalos, J. L.; Fink, G. R.; Stephanopoulos, G. Nature Biotechnology 2013, 31, 335.

(55) Zhou, Y. J.; Buijs, N. A.; Zhu, Z.; Gómez, D. O.; Boonsombuti, A.; Siewers, V.; Nielsen, J. Journal of the American Chemical Society 2016, 138, 15368.

(56) Bai, Y.; Yin, H.; Bi, H.; Zhuang, Y.; Liu, T.; Ma, Y. Metabolic Engineering 2016, 35, 138.

(57) Ahmadi, M. K.; Fang, L.; Moscatello, N.; Pfeifer, B. A. Metabolic Engineering 2016, 38, 382.

(58) Gosset, G. Current Opinion in Biotechnology 2009, 20, 651.

(59) Sun, X.; Lin, Y.; Yuan, Q.; Yan, Y. Acs Synthetic Biology 2015, 4, 554.

(60) Kunjapur, A. M.; Tarasova, Y.; Prather, K. L. Journal of the American Chemical Society 2014, 136, 11644.

